# A workflow for targeted proteomics assay development using a versatile linear ion trap

**DOI:** 10.1101/2024.05.31.596891

**Authors:** Ariana E. Shannon, Rachael N. Teodorescu, Nojoon Soon, Lilian R. Heil, Cristina C. Jacob, Philip M. Remes, Mark P. Rubinstein, Brian C. Searle

**Affiliations:** Ohio State University, Columbus, OH, United States; Pelotonia Institute for Immuno-Oncology, The Ohio State University Comprehensive Cancer Center, Columbus, OH, 43210, USA; Department of Biomedical Informatics, The Ohio State University Medical Center, Columbus, OH, 43210, USA; Thermo Fisher Scientific, San Jose, CA, 95134, United States; Division of Medical Oncology, Department of Internal Medicine, The Ohio State University College of Medicine, Columbus, OH, 43210, USA

## Abstract

Advances in proteomics and mass spectrometry have enabled the study of limited cell populations, such as single-cell proteomics, where high-mass accuracy instruments are typically required. While triple quadrupoles offer fast and sensitive nominal resolution measurements, these instruments are effectively limited to targeted proteomics. Linear ion traps (LITs) offer a versatile, cost-effective alternative capable of both targeted and global proteomics. We demonstrate a workflow using a newly released, hybrid quadrupole-LIT instrument for developing targeted proteomics assays from global data-independent acquisition (DIA) measurements without needing high-mass accuracy. Gas-phase fraction-based DIA enables rapid target library generation in the same background chemical matrix as each quantitative injection. Using a new software tool embedded within EncyclopeDIA for scheduling parallel reaction monitoring assays, we show consistent quantification across three orders of magnitude of input material. Using this approach, we demonstrate measuring peptide quantitative linearity down to 25x dilution in a background of only a 1 ng proteome without requiring stable isotope labeled standards. At 1 ng total protein on column, we found clear consistency between immune cell populations measured using flow cytometry and immune markers measured using LIT-based proteomics. We believe hybrid quadrupole-LIT instruments represent an economic solution to democratizing mass spectrometry in a wide variety of laboratory settings.

## Introduction

Systems biology is the study of interactions within and between cells, where the goal is to learn how those interactions give rise to the complex behavior seen in the entire system.^1^ One challenge is that many complex biological processes, such as adaptive immunity, are built from small populations of distinct cell types acting in concert.^2,3^ Improvements in proteomics methods and mass spectrometry (MS) instrumentation have paved the way for low-input and single-cell proteomics, which make it possible to study how limited cell populations contribute to the whole. While the majority of single-cell methods use tandem mass tags (TMT)^4^ to increase signal (and thus consistency) with data-dependent acquisition (DDA),^5,6^ several groups have demonstrated that data-independent acquisition (DIA) is an effective solution to measuring low-input samples.^7–9^ However, in nearly all cases, high-mass accuracy instruments are required.

While global proteomics on a high-mass accuracy instrument is typically used for single-cell and low-input proteomics, nominal-mass instruments, such as triple quadrupoles, have led in measurement sensitivity using targeted selection reaction monitoring (SRM).^10^ With SRM, peptides are detected based on monitoring multiple fragment ion signals produced by each selected precursor ion. Transitions (diagnostic precursor/fragment ion pairs) in a pre-specified list must be provided to the instrument for monitoring at specific times within the chromatographic gradient for a scheduled SRM.^11^ While triple quadrupoles are extremely quick instruments, capable of rapidly switching between ion pairs, they can only monitor a single *m/z* at a time. As such, triple quadrupoles are limited to targeted experiments and require a high-mass resolution instrument to select targeted peptides and transitions before migrating to a nominal-mass instrument for high-throughput monitoring.

An alternative targeted method to SRM is parallel reaction monitoring (PRM), which uses a quadrupole-equipped high-resolution mass spectrometer where the third quadrupole is replaced with a Thermo Scientific™ Orbitrap™ mass analyzer (Q-Orbitrap, also known as a Q-Exactive™) or a time-of-flight analyzer (Q-ToF). Rather than measure precursor/fragment transitions, all precursor-specific fragment ions are collected in a full tandem mass spectrum with PRM.^12^ A major advantage of PRM is that diagnostic fragment ions can be selected after an experiment through data analysis. This can vastly simplify the assay development process compared to SRM. PRM has provided meaningful biological insight into several diseases, including systemic autoimmune diseases,^13^ multiple sclerosis,^14^ and colorectal cancer.^15^ When coupled with global proteomics, PRM is a powerful tool for interrogating system-wide interactions between cells.

Linear ion traps (LITs) are another type of versatile, fast, nominal-mass analyzer, comparable in resolution and complexity to triple quadrupoles. Modern Tribrid instruments have incorporated LITs as a tertiary analyzer, coupled with an Orbitrap.^16^ Using a Tribrid™ instrument, Heil et al^17^ showed that the benefit of PRM lies within its ability to monitor multiple product ions produced within a selected precursor *m/z* range and that the LIT in Tribrids was an effective readout for targeted proteomics. A LIT measures ions trapped in an electric field by adjusting RF and DC voltages to selectively eject ions based on their *m/z* to collect MS/MS spectra. Unlike triple quadrupoles, which have to “dwell” at each increment of m/z to form a spectrum, LITs are capable of acquiring full scan MSn data in a fast and sensitive manner,^18^ making them also viable for global proteomics using DDA or DIA.^19^ As a result, a hybrid quadrupole-LIT (Q-LIT) could act as an “all-in-one” nominal-mass instrument capable of both targeted and global proteomics.

Similar to triple quadrupoles, LITs are extremely sensitive, ion-efficient mass analyzers apt for low-input proteomics.^20^ In some circumstances, LITs can be more effective than high-resolution mass analyzers for low-input samples (≤10 ng)^21^ and are capable of measuring single cells without multiplexing reagents.^22^ At higher sample input (≥100 ng) the sensitivity of LITs is overshadowed by the lack of high mass accuracy. There exist other compelling reasons to consider LIT-based instruments in high-throughput applications. In particular, LITs operate at high pressure (10^−3^ mTorr) in comparison to ToF analyzers (10^−6^ mTorr) where ions have to travel uninterrupted for meters, or Orbitrap analyzers (10^−10^ mTorr) where ions can travel for more than a kilometer. Lower vacuum pump requirements allow LITs to be built more affordably, robustly, and housed in smaller instrument footprints.

Here, we present a workflow using a hybrid quadrupole-LIT (Q-LIT) instrument from Thermo Scientific as a single instrument for rapidly generating targeted assays for low-input experiments. With the Q-LIT, we demonstrate how to generate nominal-mass targeted transition libraries using both DDA, composed of offline fractionated samples, as well as gas-phase fractionated (GPF) DIA libraries. We then show the quantitative accuracy of targeted PRMs utilizing a Q-LIT using matched-matrix calibration curves at 1, 10, and 100 ng of low abundant immune cell populations. To facilitate this, we developed an open-source software tool that directly schedule-optimized PRM assays from DIA libraries. Finally, we show the quantitative consistency measuring “real-world” biological targets in cytokine-stimulated CD8+ T cells with as little as 1 ng on column. These results suggest that a Q-LIT can perform as an inexpensive stand-alone instrument for quantitative proteomics, capable of a wide range of proteomics measurements without the need for high-resolution mass spectrometry.

## Experimental

### Splenocyte isolation and cell culture

Two murine spleens were harvested from C57BL/6J mice, counted with a hemocytometer, and then plated with anti-mCD3e (Clone 145-2C11) stimulation on the first day. This time point is referred to as Day 0. Cells were plated at a density of 3 million cells/mL in RPMI-1640, 1% non-essential amino acids + 1% penicillin and streptomycin, 1% sodium pyruvate, 0.5% β-mercaptoethanol, and 10% FBS + 0.1% and cultured at 37°C with 5% CO_2_. On Day 2, cells were washed three times with PBS, with centrifugation in between each wash at 500 RCF for 5 minutes, then replated in 24-well plates containing either human IL-2 recombinant protein or IL-15 recombinant protein at a concentration of 200 ng/mL. A third condition was maintained without stimulation as a control for flow cytometry experiments. On day 4, cells were washed and split. On day 5 of culture, one representative well from each condition for flow-activated cell sorting (FACS) using a Cytek Aurora™ flow cytometer. On day 6, a subset of cells was washed with PBS 3 times; then, cell pellets were stored at -80°C for mass spectrometry. The remaining cells were split and a representative well was again taken for FACS. On Day 10, a subset of cells was collected for flow cytometry and the remaining cells were pelleted and stored at - 80°C for mass spectrometry.

### Flow cytometry

On days 5, 6, and 10, flow cytometry was performed with the following procedure: cultures pooled together, washed three times with PBS, and counted on a hemocytometer. Cells were then stained in the dark for 30 minutes at 4°C with the viability dye (Live/Dead Fixable Blue Dead Cell Stain kit, for UV excitation from Thermo Fisher Scientific) at a ratio of 1:1000. Extracellular markers, anti-CD4, anti-CD45R, anti-CD8, and anti-TCR-β at a concentration of 1 to 400 antibody to cell solution. Additionally, anti-CD69, anti-CD62L, and anti-CD25 were added at a 1 to 200 ratio. Anti-CD44 was applied at a ratio of 1 to 600, and the cell solution was 20 minutes at 4°C. More details on the antibody panel are available in **Supplemental Table 1**. Stained cells were analyzed with a Cytek Biosciences Aurora 5-laser flow cytometer. Data was processed and visualized in BD Biosciences FlowJo™ software.

### Proteomics sample preparation

Frozen cell pellets were lysed in a 5% SDS buffer with 50 mM TEAB, 1x HALT, and 2 mM MgCl_2_. DNA was sheared with a Bioruptor® Pico by sonicating at 14°C for 30 seconds, followed by 30 seconds of rest, a total of 10 times. Sheared cells were then spun down at 13,000 RCF for 10 minutes and the protein supernatant was retained. Protein quantities were estimated using a Pierce™ bicinchoninic acid (BCA) Protein Assay Kit. Proteins were reduced with 40 mM dithiothreitol (DTT), alkylated with 40 mM iodoacetamide, and quenched with 20 mM DTT. Acidification was done with 2.5% phosphoric acid, and protein was loaded onto suspension trap (s-trap) micros (Protifi LLC). Digestion was performed with trypsin at a 1:20 ratio of enzyme to protein at 47°C for 2 hours, then eluted. Peptides were dried down and stored at -80°C.

According to the kit protocol, an aliquot of dried peptides was separated according to basicity using a Pierce High pH Reverse-Phase Fractionation Kit. Briefly, 50 μg of peptides were resuspended in 0.1% trifluoroacetic acid in HPLC-grade water. The separation mini-columns from the kit were centrifuged at 5000 RCF for 2 minutes to remove any liquid and pack the resin. The mini-columns were then equilibrated with 100% acetonitrile, and washed 3 times with water. Resuspended peptides were loaded, and the flow through was collected as the first fraction. The mini-columns were washed with water, and the eluent was collected as the second fraction. The elution buffers specified from the kit were then used to produce the following 8 fractions. Fractionated peptides were then dried down and stored at -80°C until mass spectrometry-based analysis for DDA-based library generation.

A separate aliquot of the eluted peptides was dimethyl labeled using an in-solution amine-labeling reaction published by Boresema et al.^23^ Digested peptides were resuspended in 100 mM TEAB (pH = 8.5). Formaldehyde (4%) was added to the resuspended peptides and mixed. Sodium cyanoborohydride (0.6 M) was then added to catalyze the dimethyl labeling reaction for 90 minutes at 22°C while mixing vigorously. The reaction was quenched with 1% ammonia and 5% formic acid. All peptides were resuspended in 2% acetonitrile with 0.1% formic acid. Calibration curves were generated by mixing labeled and unlabeled peptides at different concentrations. In these mixtures, unlabeled peptides were diluted in a dimethyl-labeled background over 4 orders of magnitude (**Supplemental Table 2**) and aliquoted at different concentrations prior to mass spectrometry analysis.

### LC-MS settings

Data was acquired on a Thermo Scientific™ Stellar™ MS coupled to a Vanquish™ Neo UHPLC system. Solvent A consisted of 100% water with 0.1% formic acid, and solvent B contained 80% acetonitrile with 0.1% formic acid. An Easy-Spray™ source was used for ionization at 2000 V, and the ion transfer tube was set to 275°C. Peptides were separated on a 25 cm C18 analytical Easy-Spray column, packed with 2 μm beads along a 50-minute linear gradient as follows: from 0-4 minutes, 2% B, 4-8 minutes increased to 8% B, 8 to 58 minutes increased with 28% B, 58 to 65 minutes increased to 44% B, followed by a 10-minute wash at 100% B. The flow for the entire gradient was set to 250 nL/min. The instrument was configured to expect chromatography of approximately 15 seconds, and fragment peptides with a default charge state of 2 and a collision cell gas pressure of 8 mTorr.

### DDA on an ion trap instrument

For DDA experiments, the RF lens was set to 30%. Precursor spectra were collected ranging from 350-1250 *m/z* at a scan rate of 67 kDA/s. The automatic gain control (AGC) target was set to “Standard” with an absolute AGC target of 3e4. The maximum ion injection time (maxIIT) was set to 100 ms and spectra were collected using centroiding in positive mode. MS2 scans were collected only on peptides with a charge state greater than 1, excluding undetermined charge states. An intensity threshold of 5E2 was used to trigger an MS2 scan and an HCD NCE of 30%. Following the MS2 measurement, the peptide m/zs were placed on a dynamic exclusion list for fragmentation for 2 seconds using a precursor mass tolerance of +/- 0.5 *m/z*. Twenty DDA scans were taken in each cycle with a 1.6 m/z isolation window around the precursor of interest. Fragment ions were scanned at 125 kDa/second scan rate from 200-1500 *m/z* using an AGC target of 1E4 and a maxIIT of 50 ms.

### DIA and PRM on an ion trap instrument

For both DIA and PRM experiments, the precursor range was set to 350-1250 *m/z* and measured at a rate of 67 kDa/second. The AGC target was set to “Standard,” which is equivalent to 1e4, and the maxIIT was set to 100 ms. The loop control was set to “all.” Peptides were fragmented with HCD with NCE set to 30%, and fragments were scanned at 67 kDa/second over a range of 200-1500 *m/z*. Precursor isolation windows for DIA were consistently 8 *m/z* wide, where margins were set to forbidden zone locations. Six gas phase fractions were used to collect chromatogram libraries over 400-500, 500-600, 600-700, 700-800, 800-900 and 900-1000 *m/z*. The majority of settings were the same as a wide-window DIA scan, with the exception of 2 *m/z* wide isolation windows over the adjusted precursor *m/z* range for each method.

All settings for PRM scans were the same except for the isolation window width and maxIIT. For PRM assays at 10, 20, and 50 peptides per cycle, the maxIIT were respectively set to 200, 95, and 50 ms. Precursor isolation windows were set to 2 *m/z*, where MS/MS were collected over 200-1500 *m/z*.

### Data analysis

Global data was first converted to the universal mzML format using peak picking. DIA data was analyzed with EncyclopeDIA v.3.0.0-SNAPSHOT using the Ion Trap/Ion Trap mode. Mass tolerance was set to 0.4 Da where a minimum of 3, but a maximum of 5 ions were used for quantification. The chromatogram library was generated by searching 6 gas phase fractions against a Prosit^24^ predicted library. The Prosit library contained spectrum predictions of all +2 and +3 ions from a mouse FASTA from UniProt, which was accessed on October 22, 2019. The predicted library allowed for up to 1 missed cleavage, with a default charge state of 3, and default NCE of 33 over 396.4-1002.7 *m/z*. Wide-window (8 *m/z)* injections were searched against the chromatogram library using the same search settings. Global DDA data was searched in Scribe using the Ion Trap/Ion Trap instrument mode for b and y tryptic peptides, with a library mass tolerance of 0.4 Da. The 10 high-pH fractionated injections were searched against the same Prosit predicted library used to generate the DIA library.

Global DIA data was searched using CHIMERYS^25^ intelligent search algorithm (MSAID GmbH) in Thermo Scientific™ Proteome Discoverer™ 3.1, using analogous settings for the EncyclopeDIA search. A predicted spectrum library was generated from the mouse fasta database by INFERYS™ deep learning framework (MSAID GmbH) for all tryptic +2, +3, and +4 peptides between 7-30 amino acids in length. For processing, spectrum files were selected using the ion trap MS setting, with a signal-to-noise peak threshold of 1.5. The top 24 peaks were selected in each window with a fragment mass tolerance set to 0.4 Da. Fixed carbamidomethyl modifications and a maximum of 2 oxidized methionines were allowed per peptide. The retention times from CHIMERYS were extracted for all detections and combined with EncyclopeDIA’s detections. For peptides detected by both software tools, the EncyclopeDIA retention times were preferred. The combined detections and retention times were used to select peaks within EncyclopeDIA and run against a 1% FDR to obtain a combined search engine library. The fractionated injections were also searched in Proteome Discoverer, using a mass tolerance of 0.4 Da for all +2, +3, and +4 peptides, and the same settings used for the CHIMERYS search,

Skyline^26,27^ version 23.1.0.455 was used for targeted analysis. For analyzing the calibration curves and other PRM injections of IL-2 and IL-15 replicates, the chromatogram library was first imported to serve as a reference point for integrating low-input PRMs. With the imported DIA results, transition settings were altered, and the PRM samples were imported. For both imports, the settings peptide settings were set to Trypsin [KR\P], with a maximum of 2 missed cleavage, and the mouse fasta used to generate the Prosit library was used to generate a background proteome. Retention time window predictions were set to 5 minutes, however, measured retention times were used when present. Peptides between 7 and 40 amino acids in length were used, and “auto-select all matching peptides” was left checked. Only carbamidomethylation modifications were considered for cysteine. For transition settings, peptides of +2, +3, and +4 precursor charges, along with +1 and +2 ion charges were considered for b, y, and precursor ion types. For product ion selection, we considered the third ion to the second to last ion. DIA precursor windows were used for exclusion when importing the chromatogram library. The ion match tolerance for the library was set to 0.4 Da, and 6-9 product ions were used from filtered product ions. For the instrument parameters, a 200-1500 *m/z* range was considered, with a method match tolerance of 0.4 Da, and “dynamic min product *m/z*” and “triggered chromatogram acquisition” were checked. For the full-scan parameters, DIA was used when importing the chromatogram library file as a reference point for integrating calibration curves. The gas-phase fractionated isolation windowing scheme was imported from the files for a QIT mass analyzer, with a resolution of 0.4 Da, and retention time filtering within 5 minutes of MS/MS IDs. For importing PRM injections, the “PRM” acquisition method was set, rather than the DIA method. Finally, lower limits of detection (LoD) and quantification (LoQ) were estimated using EncyclopeDIA, and calculated quantities of IL-2 and IL-15 replicates were determined using calibration curves.

### Data availability

Proteomics data and Skyline documents are available on Panorama at https://panoramaweb.org/StellarIonTrapForLowInput.url. All proteomics raw data is also publicly available on the MASSIVE repository under the accesion number MSV000094904 (ftp://massive.ucsd.edu/v08/MSV000094904/). Open-source software developed for this project is publicly available as part of the EncyclopeDIA project at https://bitbucket.org/searleb/encyclopedia.

## Results and Discussion

Linear ion traps (LITs) are robust, sensitive, fast mass analyzers, yet these instruments have limited mass resolution. Previously, our lab demonstrated that LITs could effectively be used as stand-alone mass analyzers to measure low-input samples down to single cells using a Thermo Scientific™ Orbitrap Eclipse™ Tribrid™ mass spectrometer.^22^ In that work, we detected approximately 400 proteins from single cells using data-independent acquisition coupled with chromatogram library generation to help make detections.^28^ While the Eclipse instrument configuration ignored the high-resolution Orbitrap mass analyzer, we performed those experiments in the context of a high-end Tribrid instrument. Furthermore, the 400 proteins we measured were the easiest to observe, but not necessarily the most biologically useful to monitor. While reduced representation approaches^29,30^ that monitor a limited panel of easily observed proteins can help infer biological states, significant hurdles must be overcome to predict the expression of unmeasured proteins. As such, directly measuring proteins of interest in low-input samples using targeted proteomics may be preferable to global proteomics.

In this work, we sought to answer three remaining questions. First, by eliminating the Orbitrap, could an affordable quadrupole-LIT (Q-LIT) mass spectrometer perform at a high level as a stand-alone instrument for both library generation and targeted proteomics measurement? Second, can a Q-LIT mass spectrometer quantify peptides at and below the level of single cells? Third, can low-level biologically relevant proteins be measured at or below 1 ng in quantitative experiments? To this end, we assessed several parameters of the Stellar mass spectrometer, a new hybrid Q-LIT design produced by Thermo Fisher Scientific. First, we tested proteome-wide library generation, then we assessed quantitative linearity using targeted PRM experiments with 100, 10, and 1 ng sample inputs. Finally, we tested sensitivity and measurement consistency in a biological context.

### A workflow for generating PRM assays using Q-LIT GPF-DIA data

The Stellar MS is a hybrid Q-LIT mass spectrometer with improved ion transmission features capable of performing rapid scans up to 200 kDa/s (**Figure 1A**). The instrument shares many of the same design components as existing Orbitrap-based instruments.^31–33^ Tribrid instruments accumulate fragment ions in the collision cell (also known as the ion routing multipole, IRM) before transfer to the mass analyzer. In an analogous fashion, the Stellar accumulates fragment ions in the collision cell (Q2, also known as the ion concentrating routing multipole, ICRM) prior to transfer to the LIT for mass analysis. These arrangements produce high scanning speeds by performing fragment accumulation in parallel with mass analysis in the low-pressure LIT.^34,35^

**Figure 1:**
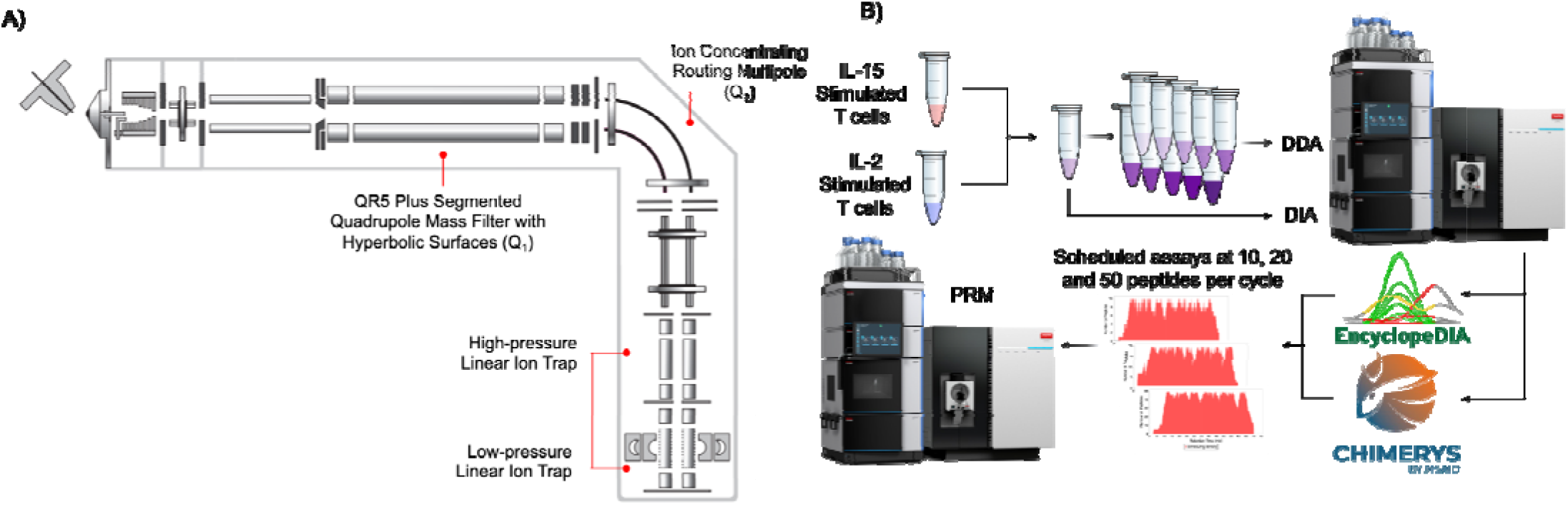
Overview of the instrument and workflow for developing targeted assays using a LIT. **A**) The instrument schematic of the Stellar MS. Ions enter the first Q5 Plus Segmented Quadrupole Mass Filter with Hyperbolic surface prior to entering into the Ion Concentrating Routing Multipole. The Ion Concentrating Routing Multipole behaves as the collision and storage cell. Ions are then moved to the high-pressure cell of the dual-pressure LIT and eventually to the low-pressure cell for mass analysis. External ion detection is performed with the Extended Scale Detector, which provides sensitive detection as an electron-multiplying device. **B**) A schematic of the methodology taken. Chromatogram libraries were generated using a GPF-DIA approach, and DDA libraries were generated from offline high-pH reverse-phase fractionated proteomes. Samples were searched by both EncyclopeDIA and CHIMERYS, and the combined results were used to schedule PRM assays at the 100, 10, and 1 ng levels.

An advantage of the Q-LIT geometry is that it is suitable for both global discovery proteomics as well as targeted proteomics. To leverage this, we implemented a workflow to generate high-quality peptide libraries using off-line fractionated DDA or GPF-DIA, and software to build on-the-fly PRM assays for the same instrument (**Figure 1B**). Briefly, the software takes the DDA-based spectrum library or a DIA-based chromatogram library as input and aligns that against potential targeted proteins. The target list contains a list of desired accession numbers along with other optional proteins in a selected FASTA database. The assay can be modified using both a peptide inclusion and exclusion list. Assays can be adjusted depending on instrument settings, where the maximum assay density and a retention time scheduling window width must be selected. Additionally, a recent single-injection DIA run ensures that retention time scheduling matches the current LC column conditions.

Peptides are selected using this software tool based on the highest signal recorded in the library. If DIA data was used for library generation, then that signal is based on the intensity of the third largest fragment ion for that peptide, following common SRM conventions of requiring at least three transitions.^36^ Peptides are chosen using a greedy approach, where the most abundant peptides are scheduled first. After a specified number of peptides are chosen for a given protein, no additional peptides from that protein are added.

Additionally, peptides cannot be added to a retention time region if any time point in that region has already reached the maximum assay density. Once all possible peptides are considered, the software tool produces a scheduling report as well as a target inclusion list for the Thermo method editor. This software workflow has a graphical user interface built into the EncyclopeDIA code base.

### Combining search engine results to develop a comprehensive library

PRM assays are commonly generated from various sources, including public repositories that store targeted proteomics, such as the PeptideAtlas,^37^ CPTAC,^38^ or Panorama^39^. Additionally, assays can be built from a global experiment, with targets selected from empirical data on the biological matrix of interest. For this work, we wanted to use methods that could be fully acquired on the Q-LIT, but still be capable of detecting low abundant peptides. One advantage of this approach is that target chemical data is tuned for the PRM instrument from the context of retention time scheduling and optimal transition selection.

To generate a PRM assay for low input on the Q-LIT, we tested two common methods of building libraries: a chromatogram library using GPF-DIA and a spectral library using fractionated DDA. Both libraries were collected from a pool of IL-2 and IL-15-stimulated T cell proteomes. To build the chromatogram library, 6x gas-phase fractions were used with 2 *m/z* wide isolation windows across mass ranges of 100 *m/z* per injection. The DDA was fractionated offline using high-pH reverse phase separations to yield a total of 10 fractions. The DDA and DIA libraries shared around half of the total peptides detected (**Figure 2A**), presumably due to the different fractionation methods used to generate each library. For this work, peptides in each DIA library were fractionated in the gas phase, where the proteome is not chemically altered or diluted. As a result, this approach produces retention times that more closely match the quantitative PRM experiments, since the matrix background remains consistent. In contrast, peptides in each DDA library are chemically fractionated by reverse-phase chromatography. Consequently, each fraction has a simplified matrix background, which may not reflect retention times as consistently without similarly fractionating each quantitative sample.

**Figure 2:**
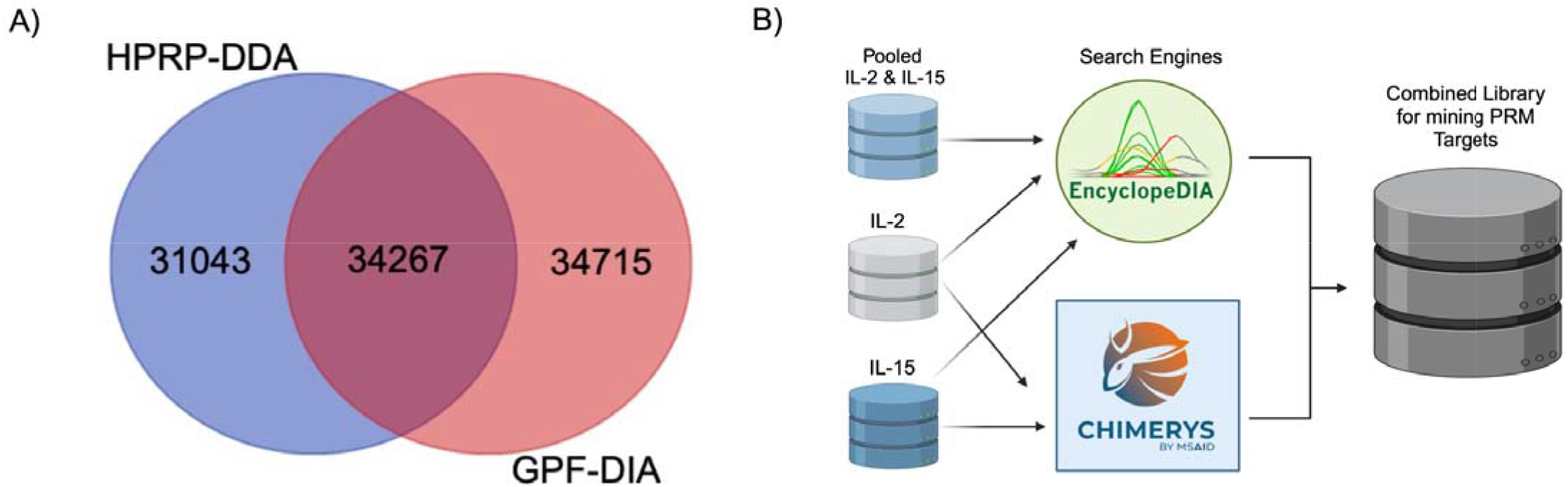
A spectral library using offline fractionated data-dependent acquisition (DDA) and the chromatogram library using data-independent acquisition (DIA). **A**) A Venn diagram of the peptides detected from each library. **B**) Detections from CHIMERYS were combined with EncyclopeDIA to generate a combined search engine library to mine PRM targets.

In addition to producing more peptide detections, peptide-centric extraction^40^ of DIA datasets is more akin to fragment-level quantification using targeted methods than DDA measurements.^41^ As such, peptides detected using GPF-DIA are more likely to produce robust targeted assays since the mode of discovery uses similar methodologies to the final quantitative measurements. However, some sample types, such as enriched phosphopeptides, are better suited to library generation with DDA since they can take advantage of stochastic sampling to detect more peptides and proteins over technical replicates.^42^ For this work, we chose to proceed with the GPF-DIA library for assay development, but the scheduling software produced for this work functions with either library source.

For DIA injections, search results from EncyclopeDIA and CHIMERYS were combined for downstream work (**Figure 2B**). More peptides were detected from CHIMERYS compared to EncyclopeDIA in each gas-phase fraction, but considering the superset of detections increased the total number of potential targets (**Supplemental Figure 1**). CHIMERYS is a spectrum-centric search engine that builds on INFERYS to provide spectra and retention-time predictions for peptides in a given FASTA database.^43^ In comparison, we searched a Prosit-predicted spectral library^24,44^ with the peptide-centric search engine, EncyclopeDIA, which was adapted for analyzing ion trap data. The retention times from CHIMERYS-detected peptides were re-peak picked using EncyclopeDIA to identify candidate target transitions for PRM measurement in a combined DIA library.

### Assessing Q-LIT PRM quantitative accuracy at low input

With low-input global proteomics, we preferentially measure only the most abundant proteins. We stress-tested the quantitative accuracy of the Q-LIT system using PRMs by measuring biologically relevant proteins that tend to occur at various levels in the proteome. To accomplish this, we first functionally annotated candidate peptides in the combined DIA library using the PANTHER database.^45^ We selected target proteins based on GO-terms and Reactome pathways for T cell differentiation, immune biology, T cell activation, cytokines, and transcription factors. Using the new PRM scheduling algorithm, we constructed three assays using the same bank of proteins, where each assay was suited to a different input level: up to 50 peptides/cycle for 100 ng of material, 20 peptides/cycle for 10 ng of material, and 10 peptides/cycle for 1 ng of material. Ultimately, the 100 ng assay monitored 481 peptides, the 10 ng assay monitored 151, and the 1 ng assay monitored 61. To maintain a 2-second cycle time using 1 ng of material, the maximum ion injection time (maxIIT) was set to 200 ms. Similarly, at 10 ng of material the maxIIT was set to 95 ms (slightly below 100 ms to accommodate additional time required to route ions in the mass spectrometer). At 100 ng of material, the ion injection time was set to 50 ms; however, each scan rarely met that length of time.

We performed matrix-matched calibration curves^46^ at 100 ng, 10 ng, and 1 ng levels to assess the quantitative accuracy of the Q-LIT over several orders of magnitude. Dilutions in a buffer background are useful to assess instrument sensitivity, but because background noise decreases at the same rate as target peptides, quantitative linearity will always appear to be more accurate than in a real background matrix. Matrix-matched calibration curves are more effective at assessing linearity in real-world scenarios since the background signal does not change with dilution. To accomplish this, we had to build a suitable background matrix of similar composition to our target T cell proteome. Our approach used dimethyl labeling to modify the foreground T cell proteome, which kept the same composition while also producing different precursor and fragment masses. Dimethyl labeling was first introduced as a multiplexing method where multiple samples would be labeled and mixed prior to mass spectrometry.^23^ In our approach, only the background is modified, where free amines are mass-shifted by two methyl groups (28 Da). This shifts any labeled precursors (even with a single methyl group) outside of the precursor isolation window used by PRM measurements, ensuring that any foreground signals will not be confused with background signals. Additionally, dimethyl labeling is affordable, easy, and quick, as peptides are labeled to 99.9% completion within a 1-hour reaction.

While ion traps generally have a more limited dynamic range compared to Orbitrap-based mass spectrometers, the sensitivity of ion traps allows for superior detection and quantification of low-input samples.^17^ Additionally, Orbitraps have slower scanning speeds compared to the Q-LIT, limiting the number of peptides that can be targeted within a cycle. Here, we show that you can also achieve reasonable quantitative accuracy with the Q-LIT at low input while targeting a similar number of peptides as is possible in higher-input PRM assays with an Orbitrap. At 100 ng, the quantitative accuracy of most peptides acquired with PRM remains consistent for nearly two orders of magnitude (**Figure 3A**), where the median lower limit of detection (LoD) was 1:4.4 and the median lower limit of quantification (LoQ) was 1:2.7 (**Figure 3B**). At the 10 and 1 ng levels, the median quantitative accuracy started to drop after one order of magnitude, where 4.34% and 1.6% of peptides, at the 1 and 10 level respectively, could not be assigned an LoQ. Comparatively, only 1.2% of peptides at the 100 ng level were unable to be assigned LOQs. At the 100 ng level, signal is more easily distinguishable from noise and the LOD distribution is generally higher than at 10 ng or 1 ng.

**Figure 3:**
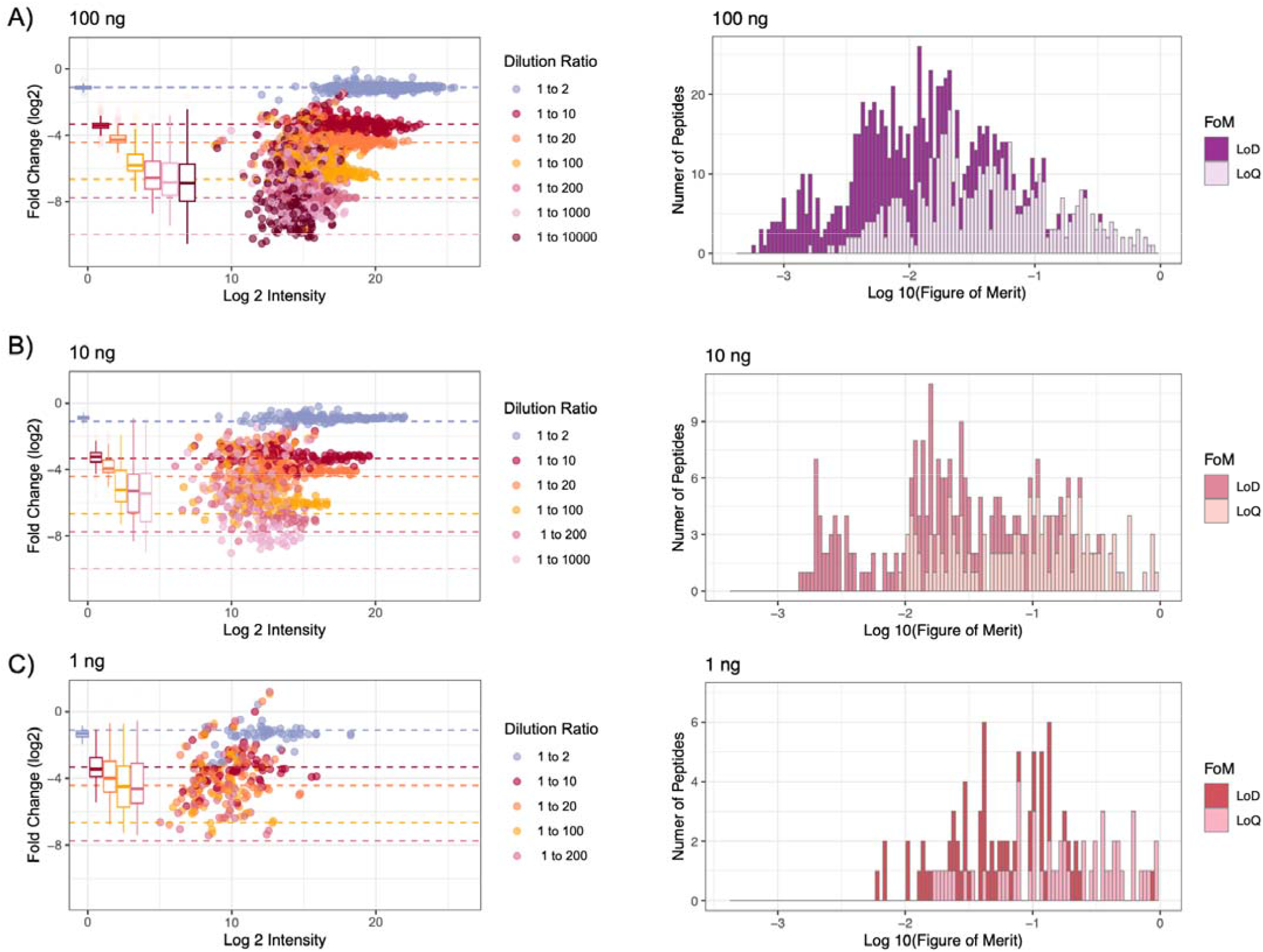
**A**) The quantitative accuracy of matrix-matched curves on an ion trap of pooled IL-2 and IL-15 peptides in a background of dimethyl-labeled pooled peptides. Three curves were generated, over 4 orders of magnitude in each, while loading 100 ng, 10 ng, and 1 ng of material on-column. Each dilution is a different color and colored dashed lines for each dilution to show where the fold change ideally should land. Box plots show the spread of measured values where the whiskers indicate 5% and 95% points and the bold line indicates the median measurement. **B**) For each curve, there is a histogram of the number of peptides with assigned lower limits of detection (LoD) and quantification (LoQ).

Single cells typically produce between 0.1 and 0.4 ng of peptides, depending on the cell type. Considering the 1 ng sample, the median measured peptide produced a linear signal in this range. While not all peptides were consistently detected at this level, several peptides showed a linear response below 0.1 ng. For example, the peptide ECESYFK from granzyme B was found to have a LoQ of 0.043:1, equating to a proteome fraction consisting of 43 pg in a background of 1 ng, and was still measurable above background an order of magnitude below that level (**Figure 4A and 4B**). Granzyme B is a serine protease implicated in multiple autoimmune diseases.^47^ All told, 61 peptides with estimated LoQs in the 1 ng assay corresponed to 30 quantified proteins. At least 6 points across the peak were sampled for all peptides in the assay, with a goal of achieving 8-10 points across the peak on average.

**Figure 4:**
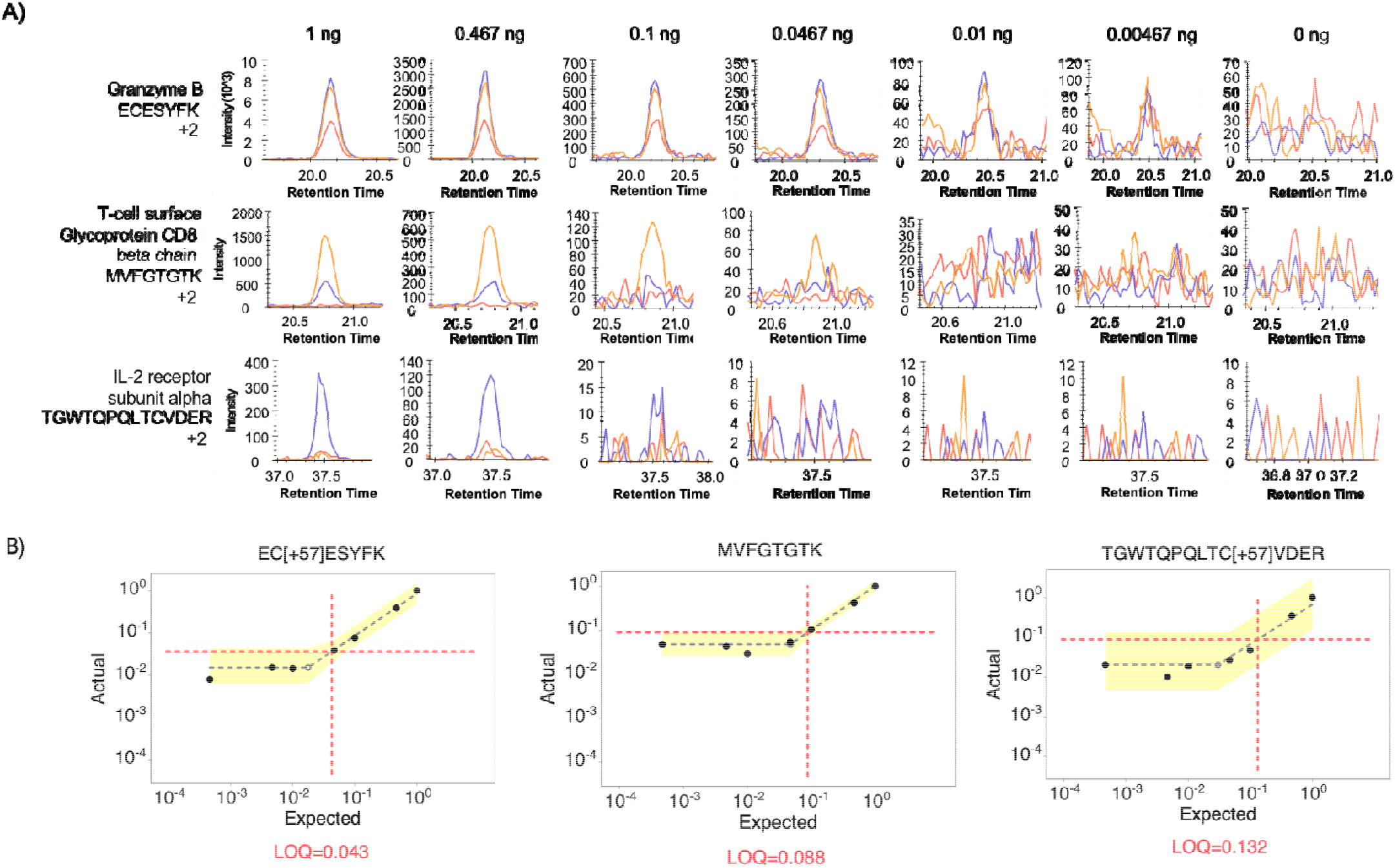
Three representative peptides that were quantifiable below 1 ng. **A**) Each row displays a peptide chromatogram at each dilution within the 1 ng curve. Each peptide contains three representative transitions. The first peptide from Granzyme B has the best estimated LoQ at 0.043, while the third peptide from IL-2 receptor subunit alpha, has the weakest estimated LoQ at 0.132 ng. **B**) LoQ and LoD were estimated on a peptide-by-peptide basis.

### Validated cell populations for experimental testing

In addition to showing quantitative accuracy in a controlled matched matrix, we wanted to validate measurement precision in low-input biological experiments. The interleukins (IL) family of proteins is a class of cytokines expressed by immune cells, including T cells, which display pro- and anti-inflammatory roles. Specific cytokines, such as IL-2 and IL-15, are essential for the proliferation and differentiation of T cells; however, each protein will enable the differentiation of divergent populations of CD4+ and CD8+ T cells. In this study, peripheral blood mononuclear cells were isolated from the spleens of 2 mice and then cultured in the presence of either IL-2 or IL-15 for up to 8 days (**Supplemental Figure 2A**). We selected this model system to showcase the ability to generate LIT-PRM assays using well-studied biology below 1 ng. Additionally, flow cytometry was used as an orthogonal technique to validate the cell populations present in IL-2 and IL-15 T cells on days 5, 6, and 10 of cell culture to measure the extent of stimulation each cytokine elicited (**Supplemental Figure 2B and 2C**).

By day 10, several populations of T cells were present in each culture, predominately CD8+ T cells in both IL-2 (83.8%) and IL-15 (92.2%) stimulated CD8+ T cells (**Figure 5**). While IL-2 is one of the first cytokines to show a response in clinical patients,^48,49^ an IL-15 analog has been shown to increase the presence of cytotoxic T cells in patients.^50^ Recently, a chimeric IL-15 analog, named “nogapendekin alfa inbakicept” has been found to act as a superagonist and has been approved as a treatment for bladder cancer.^51^ IL-2 increases the presence of regulatory CD4+ T cells, while IL-15 increases the presence of CD8+ effector cells, a cytotoxic cell population, and the basis of CAR-T therapy.^52^ We found this reflected in the cytokine stimulation assay, as CD4+ T cells composed 14.6% of the cells stimulated with IL-2. Comparatively, only 6.9% of CD4+ T cells were present in the culture stimulated by IL-15.

**Figure 5:**
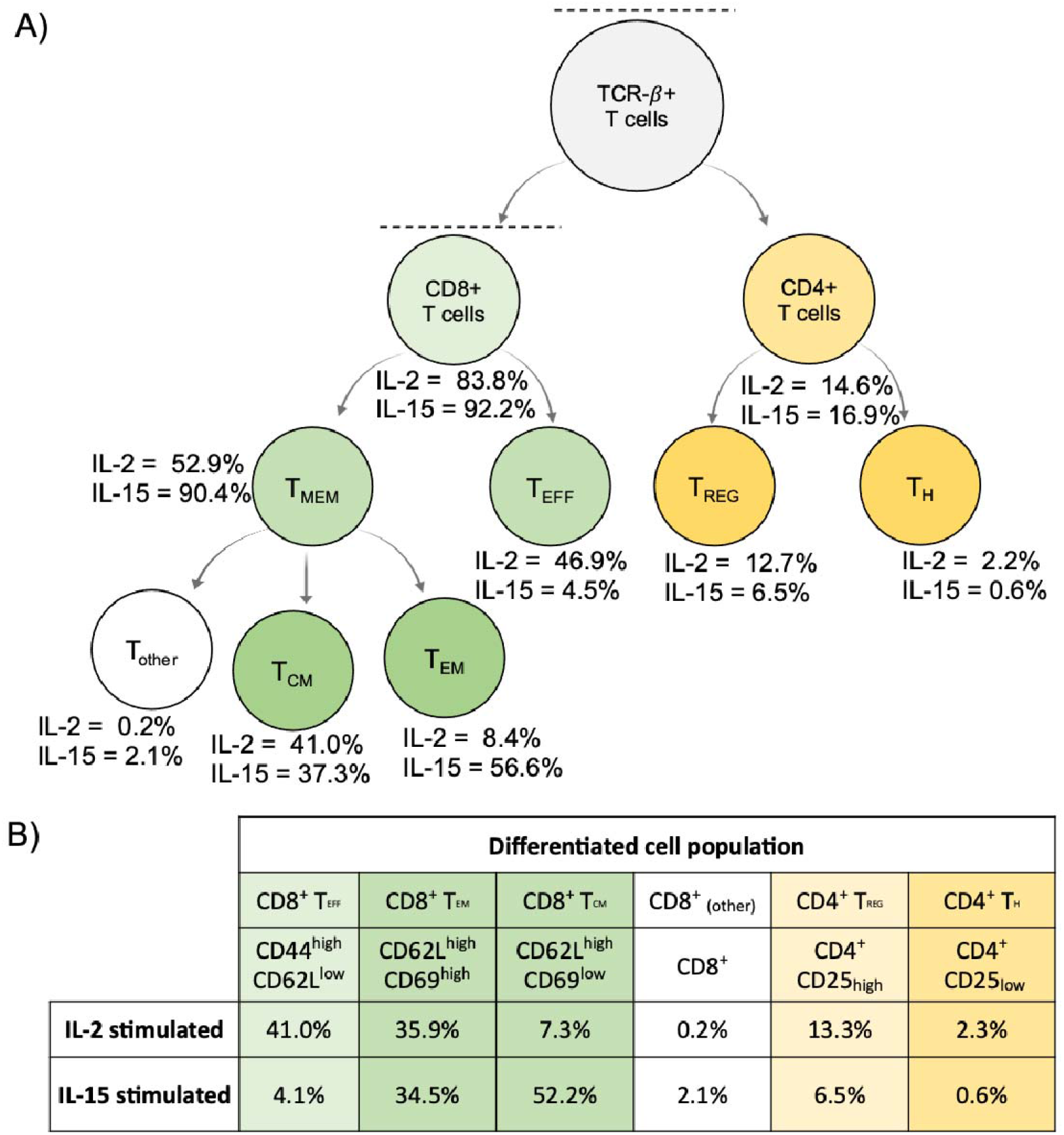
A summary of the cell populations in IL-2 and IL-15 stimulated T cells determined by the flow cytometry panel described in **Supplemental Table 1. A**) The gating procedure used for determining the relative percentage of each cell type in the IL-2 and IL-15 samples (more details in **Supplemental Figure 2**). **B**) The estimated cell populations based on back calculations of the gating results.

Using the Q-LIT, we performed 1 ng injections in technical triplicate of the IL-2 and IL-15 stimulated T cells from the final (day 10) samples. IL-2 stimulation pushes naive splenocytes to differentiate into effector CD8+ T cells, which is reflected in the flow cytometry data. Granzyme B, from which we estimated that at least one peptide was quantitative to 0.043:1 (ratio of foreground to background) is an effector molecule, secreted by cytotoxic CD8+ T cells. We found that peptides associated with this protein were 12x lower in IL-15 than IL- 2 stimulated cells (**Figure 6A**). This result is consistent with the flow cytometry data, where the effector CD8+ T cells (T_EFF_) were 10x lower in abundance when stimulated with IL-15 compared to IL-2.

**Figure 6:**
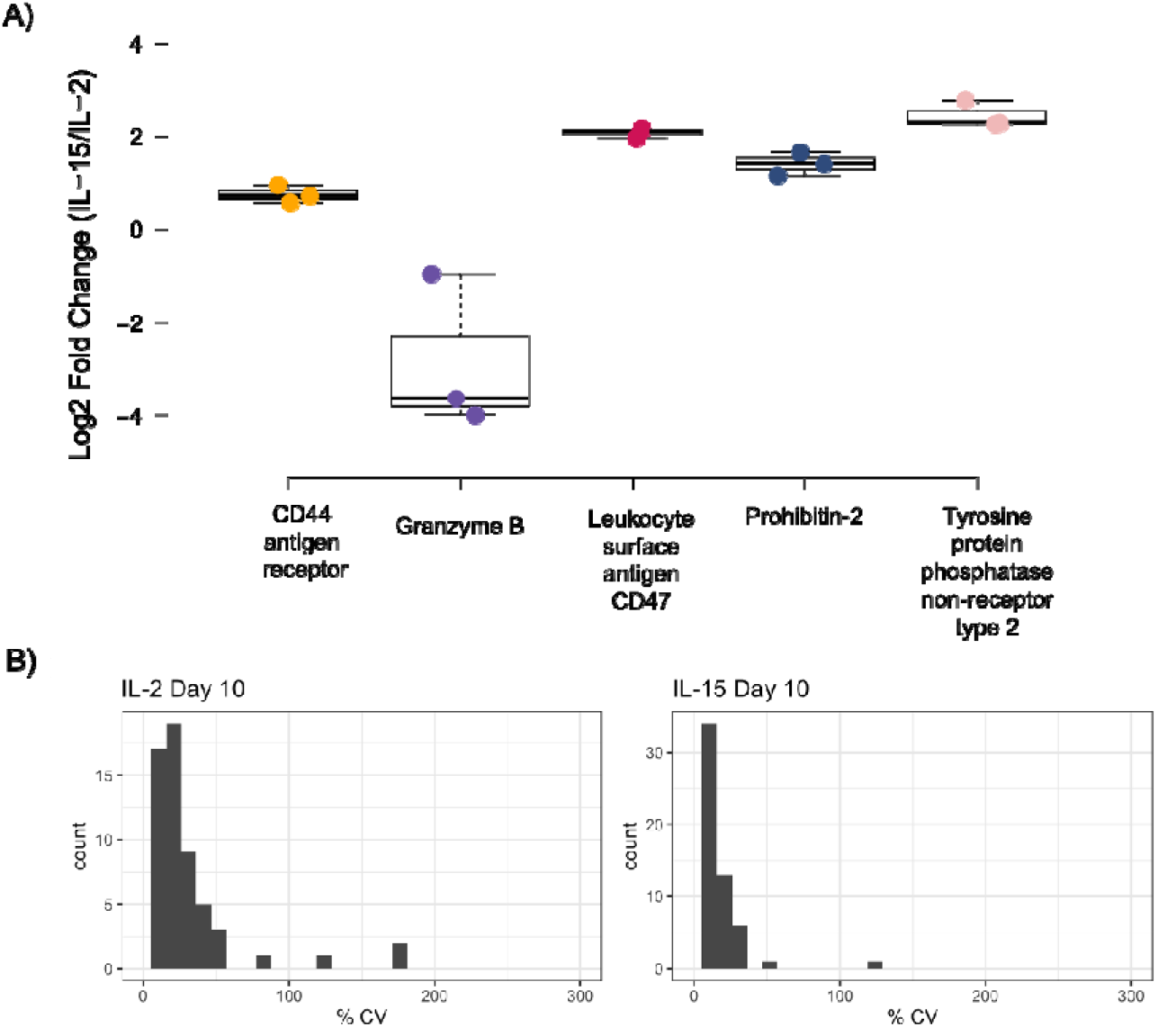
Quantifying biological replicates from the calibration curves at 1 ng. **A**) Quantitative ratios for five proteins from technical triplicate injections of IL-2 and IL-15 stimulated proteomes. **B**) Coefficient of variation (% CV) plots for all peptides quantified in the 1 ng assay.

IL-15 stimulation leads to the differentiation of memory T cells, which is demonstrated in the flow cytometry data. CD44 receptor antigen is a cell surface receptor that helps cells facilitate cell-cell interaction and respond to the tissue microenvironment. T cells that are high in this marker are found to have an increased population of memory T cells after IL-15 stimulation.^53^ Flow cytometry data indicated that we had a higher population of CD44^High^ and CD62L^High^ cells in the IL-15 stimulated condition, compared to T cells stimulated with IL-2 (**Supplemental Figure 2**). Within the PRM at the 1 ng level, we find that the expression of CD44 is slightly higher in IL-15 compared to IL-2, with a median of a 0.73 log 2-fold change in abundance. In general, we observe very high precision using the Q-LIT platform, even in 1 ng PRM assays, where most peptides fall below 20% coefficient of variation (**Figure 6B**).

Within our flow cytometry data, we tracked central memory and effector memory cells in the IL-2 and IL- 15 stimulated cultures. We found that the effector memory population was equivalent in both cell populations; however, the central memory population was greatly increased in the IL-15-stimulated cells. We are not able to see all of the markers used in the flow cytometry panel in the proteomics data at the 1 ng level, including the memory marker CD62L, yet we can use both techniques as complementary methods and show the same biological trends.

## Conclusions

Here we demonstrate a complete workflow for acquiring global libraries and rapidly producing PRM assays using only a Q-LIT mass spectrometer. While the Q-LIT is capable of DDA, we found that large libraries could be easily generated using GPF-DIA. Using the Q-LIT, we found PRM assays to be highly quantitative, even at low input. From a wider perspective, high-resolution mass spectrometry is expensive both in terms of instrument costs and in requiring higher technical experience to successfully operate. In contrast, Q-LIT mass analyzers are easier to maintain and more cost-effective to operate because they have less stringent vacuum requirements compared to Orbitrap-based mass spectrometers, making them an interesting option for low-input proteomics. These factors are especially important in single-cell proteomics, where thousands of injections are needed for each biological sample. Our results suggest that Q-LITs provide laboratories with highly capable low-input proteomics analysis in situations where high resolution is unnecessary. We believe these instruments provide a high value-to-expense ratio, potentially democratizing the use of mass spectrometry in a wider array of laboratory settings.

## Acknowledgements

We thank Zihai Li for helpful discussions. This research was supported by NIH R35GM150723 and R21CA267394 to BCS. This research was also supported by The Ohio State University Comprehensive Cancer Center (OSUCCC) and the National Institutes of Health (NIH) under grant P30CA016058. This research was made possible through resources, expertise, and support provided by the Pelotonia Institute for Immuno-Oncology (PIIO), which is funded by the Pelotonia community and the OSUCCC. We thank the PIIO and the Immune Monitoring and Discovery Platform for flow cytometry access.

## Supplementary Materials

**Supplemental Table 1**: Flow cytometry panel

**Supplemental Table 2**: Dilutions used for calibration curves at 100 ng, 10 ng, and 1 ng of total material

**Supplemental Figure 1**: Comparison of Chimerys and EncyclopeDIA detections for chromatogram libraries

**Supplemental Figure 2**: Additional flow cytometry validation data

