## Supplementary Materials for "A workflow for targeted proteomics assay development using a versatile linear ion trap"

**Supplemental Table 1**: Flow cytometry panel

| **Marker** | **Channel** | **Fluorochrome** | **Vendor** | **Clone** | **Catalog** | **Concentration** | **Staining** |
| --- | --- | --- | --- | --- | --- | --- | --- |
| **CD4** | UV2 | BUV395 | BD Biosciences | GK1.5 | 563790 | 7:40 | Extracellular |
| **Viability** | UV6 | Live/Dead Fix Blue | Thermo Fisher Scientific |  | L23105 | 17:40 | Viability |
| **CD45R (B220)** | UV10 | BUV615 | Thermo Fisher Scientific | RA3-6B2 | 366-0452-82 | 7:40 | Extracellular |
| **CD8** | UV16 | BUV805 | BD Biosciences | 53-6.7 | 612898 | 7:40 | Extracellular |
| **CD69** | V2 | SuperBright 436 | Thermo Fisher Scientific | H1.2F3 | 62-0691-82 | 4:20 | Extracellular |
| **CD44** | B3 | Alexa Fluor 532 | Thermo Fisher Scientific | IM7 | 58-0441-82 | 11:00 | Extracellular |
| **CD62L** | YG3 | PE-CF594 | BD Biosciences | MEL-14 | 562404 | 4:20 | Extracellular |
| **TCR BETA** | YG9 | PE-Cy7 | Thermo Fisher Scientific | H57-597 | 25-5961-82 | 7:40 | Extracellular |
| **CD25** | R4 | Alexa Fluro 700 | Thermo Fisher Scientific | PC61.5 | 56-0251-82 | 4:20 | Extracellular |

**Supplemental Table 2**: Dilutions used for calibration curves at 100 ng, 10 ng, and 1 ng of total material on-column.

| **% Foreground** | **% Background** | **Amount of material for the 100 ng Calibration curve** | **Amount of material for 10 ng**  **Calibration curve** | **Amount of material for 1 ng**  **Calibration curve** |
| --- | --- | --- | --- | --- |
| 100.00% | 0.00% | 100.000 | 10.000 | 1.000 |
| 46.67% | 53.33% | 46.667 | 4.667 | 0.467 |
| 10.00% | 90.00% | 10.000 | 1.000 | 0.100 |
| 4.67% | 95.33% | 4.667 | 0.467 | 0.047 |
| 1.00% | 99.00% | 1.000 | 0.100 | 0.010 |
| 0.47% | 99.53% | 0.466667 | 0.046667 | 0.004667 |
| 0.10% | 99.90% | 0.100 | 0.010 | - |
| 0.01% | 99.99% | 0.010 | - | - |

**Supplemental Figure 1**: Comparison of Chimerys and EncyclopeDIA detections for chromatogram libraries.

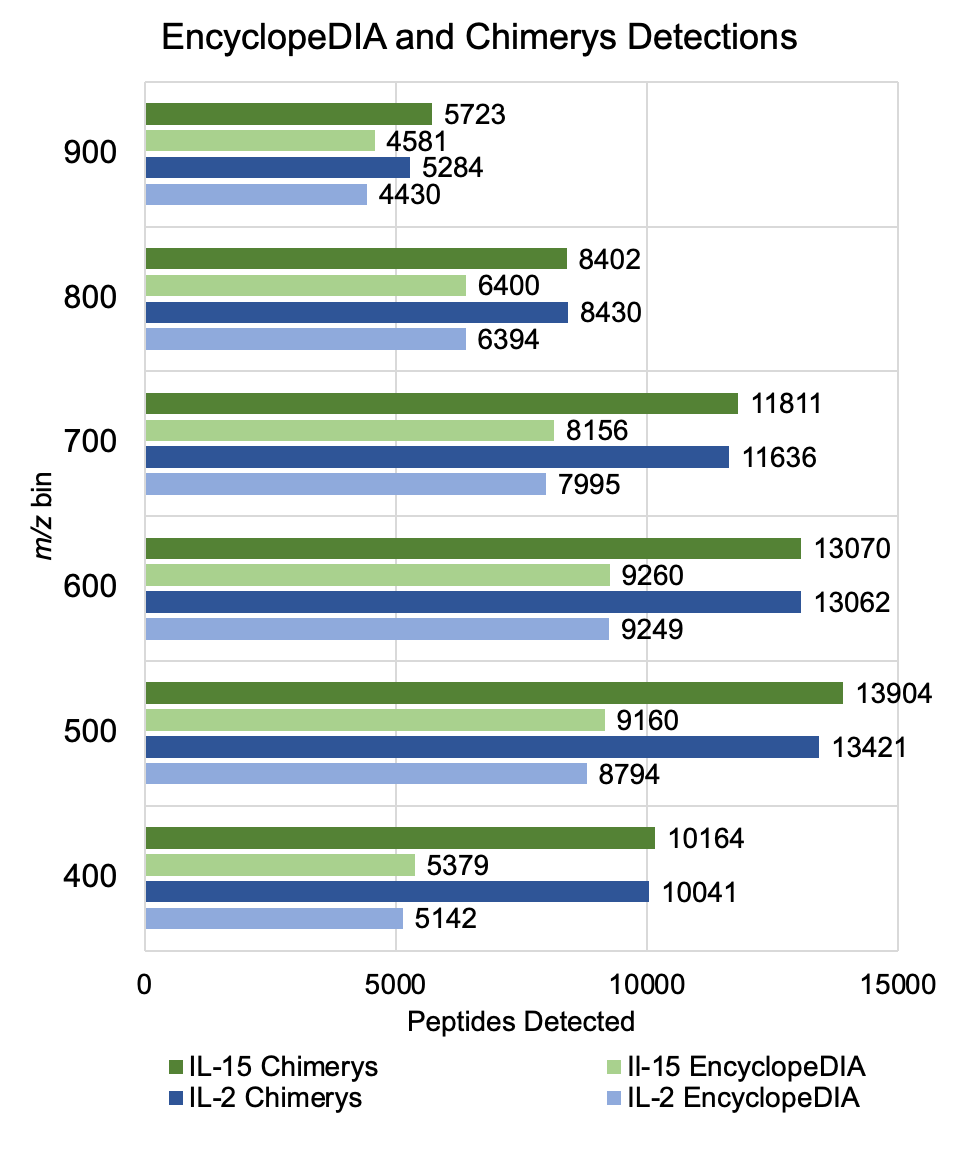

**Supplemental Figure 2**: Additional flow cytometry data

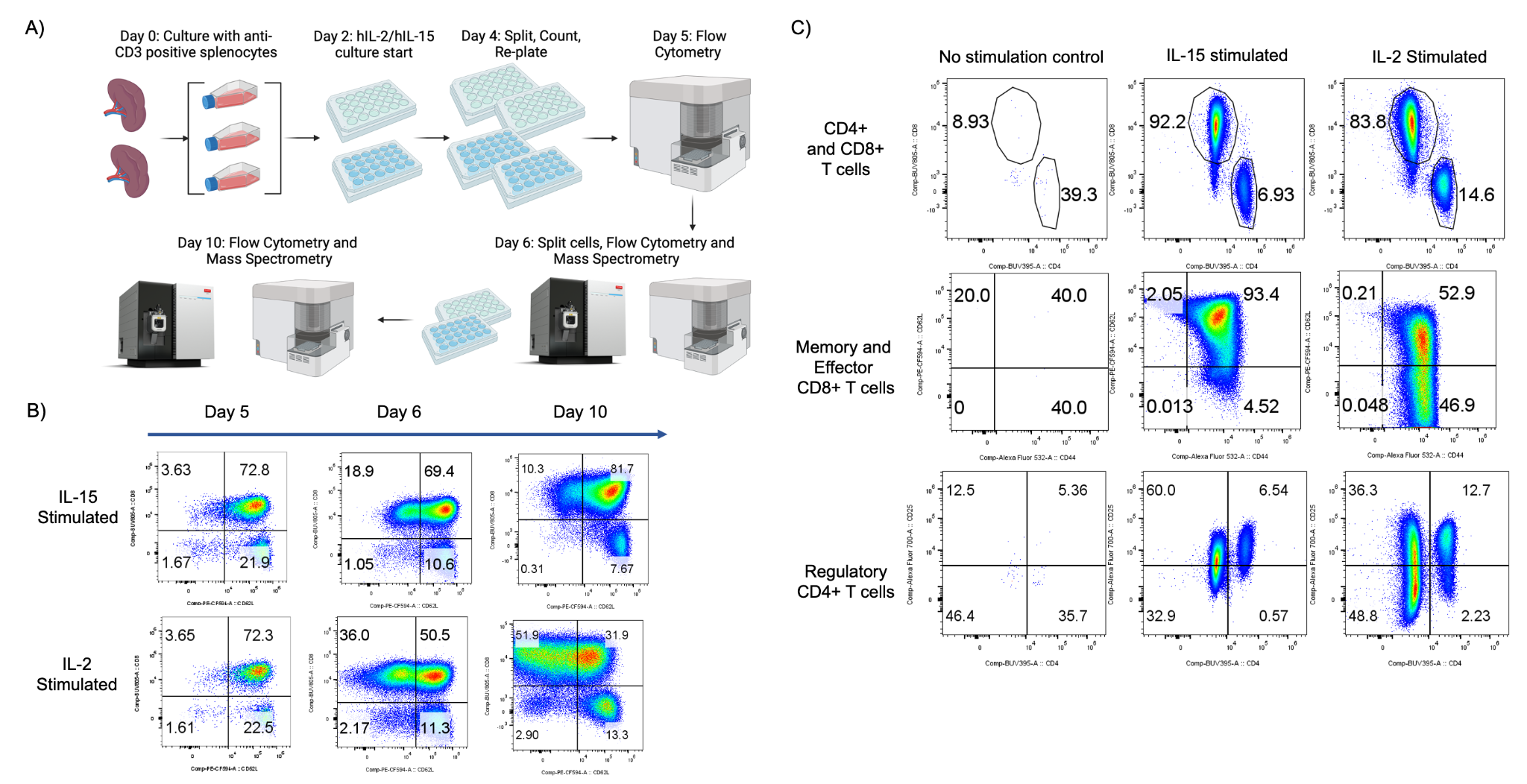
